# Estrous phase during fear extinction modulates fear relapse through a nigrostriatal dopamine pathway

**DOI:** 10.1101/2024.05.31.596838

**Authors:** Alyssa A Hohorst, Margaret K Tanner, Rebecca Han, Kamryn Korth, Jessica D Westerman, Carolina Sanchez Mendoza, Miles Q Dryden, Lareina Alverez, Remla Abdul, Esteban C Loetz, Erik B Oleson, Benjamin N Greenwood

## Abstract

Elevated ovarian hormones during fear extinction can enhance fear extinction memory retention and reduce renewal, but the mechanisms remain unknown. Ovarian hormones modulate dopamine (DA) transmission, a key player in fear extinction. In males, stimulation of substantia nigra (SN) DA neurons during fear extinction reduces renewal; an effect mimicked by a DA D1 receptor agonist into the dorsolateral striatum (DLS). The current studies tested the role of the SN-DLS pathway in estrous cycle-modulation of fear extinction and relapse. In cycling female, Long-Evans rats, fear extinction during proestrus or estrus (Pro/Est; high hormones) resulted in less relapse (renewal and spontaneous recovery) compared to males or females in metestrus or diestrus (Met/Di; low hormones). This effect was mimicked by estradiol (E2) administration to ovariectomized rats. Females in Pro/Est had greater fear extinction-induced cFos within SN DA neurons compared to males. Similarly, fast scan cyclic voltammetry revealed that electrically-evoked DA release in the DLS is potentiated by E2 and is greater during Pro/Est compared to Met/Di. An inhibitory intersectional chemogenetic approach targeting the SN-DLS pathway suppressed electrically-evoked DA release in the DLS and restored fear renewal in females exposed to simultaneous fear extinction and SN-DLS inhibition during Pro/Est. Conversely, chemogenetic stimulation of the SN-DLS pathway during extinction reduced fear renewal in males. These data suggest that levels of ovarian hormones present during fear extinction modulate relapse through a SN-DLS pathway, and that the SN-DLS pathway represents a novel target for the reduction of fear relapse in both sexes.

## Introduction

Fear extinction memories remain susceptible to fear relapse phenomena such as fear renewal (return of fear in contexts different from where extinction took place) and spontaneous recovery (return of fear after the passage of time; [1, 2]). Although means to enhance fear extinction retention have been identified, manipulations of known extinction circuits have had limited success in reducing relapse in a clinical setting. Novel strategies for the reduction of fear relapse after extinction are needed.

Biological sex is one factor known to modulate fear extinction, although recent work on sex differences in fear renewal reveal conflicting results [3, 4]. The discrepancies between studies could be due to variations in estrous phase at the time of fear extinction. Indeed, female rodents and humans exposed to fear extinction under conditions of high estradiol (E2), either via E2 administration or reproductive cycle phases during which E2 is elevated (proestrus and estrus in rodents; Pro/Est), have improved fear extinction memory compared to low E2 females (metestrus and diestrus in rodents, Met/Di; [5-12]. Similarly, Bouchet et al. (2017) reported that females exposed to fear extinction during Pro/Est have less renewal than females extinguished during Met/Di [13]. However, how levels of renewal in Pro/Est and Met/Di females compare to males, whether estrous phase during fear extinction impacts other forms of relapse, and the mechanisms involved in estrous cycle-modulation of fear extinction and relapse remain unknown.

Dopamine is emerging as a key player in fear extinction [14-16]. Importantly, augmenting fear extinction with DA agonists seems to render fear extinction memories resistant to relapse. For example, the DA agonist L-DOPA administered after fear extinction reduces the renewal and spontaneous recovery of conditioned fear in male mice and humans [17, 18].

Elevated E2, either via estrogen administration [19] or during Pro/Est [20, 21], is positively associated with potentiated stimulus-evoked DA release. It is therefore possible that high levels of ovarian hormones during fear extinction renders fear extinction memory resistant to relapse through a mechanism involving DA.

Unique roles of ascending DA pathways in fear extinction are being uncovered. The mesocorticolimbic pathway, originating in the ventral tegmental area and projecting to forebrain regions classically implicated in fear extinction is implicated in fear extinction acquisition and consolidation [14-16, 22]. However, whether alterations in DA transmission in these regions can impact fear relapse is not well understood. The nigrostriatal pathway, consisting of midbrain DA neurons arising from the substantia nigra (SN) and projecting to the dorsal striatum (DS), has been increasingly implicated in behaviors beyond its traditional role in movement [23-29], including fear extinction and relapse. For example, chemogenetic stimulation of SN DA neurons during fear extinction prevents fear renewal in male rats [30]. This is particularly interesting given that females have potentiated stimulus-evoked DA release in the DS when in Est compared to Di [20]. The dorsolateral (DLS) and dorsomedial (DMS) subregions of the DS subserve unique functions, and the effect of SN DA stimulation is mimicked by D1R agonist injected into the DLS, but not DMS, in males [31]. These data raise the possibility that SN-DLS pathway activity mediates the effects of estrous cycle on fear extinction and relapse.

The current studies investigated the role of the SN-DLS pathway in estrous cycle-modulation of fear extinction and relapse. We observe that females undergoing fear extinction in Pro/Est have less fear relapse compared to males and females in Met/Di, an effect that is replicated by E2 administration. High levels of ovarian hormones are associated with potentiated activity of the SN-DLS pathway. SN-DLS pathway inhibition during fear extinction restores relapse in females exposed to fear extinction during Pro/Est, and its activation during fear extinction reduces relapse in males. Results provide new insight into how the estrous cycle interacts with the nigrostriatal pathway to render fear extinction memory resistant to relapse and implicate the SN-DLS pathway as a novel target for the reduction of fear relapse in both sexes.

## Materials and Methods

### Animals and Housing

Fifty-eight male and 144 female (P56 on arrival) Long-Evans rats (Charles River, Wilmington, MA) were pair-housed in ventilated cages (24 L x 45.4 W x 21 H cm) in a temperature-(22° C) and humidity-controlled vivarium accredited by the Association for Assessment and Accreditation of Laboratory Animal Care located on the University of Colorado Denver Auraria campus. All experimental protocols were approved by the University of Colorado Denver Animal Care and Use Committee.

### Estrous Phase Identification

Vaginal lavages were taken and analyzed as previously described (Tanner et al. 2023; Supplementary Materials) daily for 7 consecutive days prior to start of behavioral procedures, as well as ∼2 h prior to, and immediately after each behavioral test. Males were handled for an equivalent period, if applicable.

### Surgical Procedures

All surgery was performed under Ketamine (75.0 mg/kg i.p.) and Medetomidine (0.5 mg/kg i.p.) anesthesia. Injections of carprofen (5 mg/kg s.c.) and penicillin G (22,000 IU/rat s.c.) were administered at induction and every 24 h for 72 h after surgery and rats recovered for at least 2 weeks prior to experimentation. Bilateral ovariectomy (OVX, n = 23), was performed as previously described [32]. Stereotaxic viral infusions were administered bilaterally via Hamilton needles connected to infusion pumps (World Precision Instruments, Sarasota, FL) at a rate of 0.1 μL/min. All rats received bilateral infusion of AAV2/retro-eSYN-EGFP-T2A-iCre-WPRE into the DLS (+0.5 mm anterior, ± 3.9 mm lateral, -5.4 mm ventral from the top of the skull). Rats in control groups received AAV8-hsyn-DIO-mCherry (mCherry; Addgene) into the SN (−5.4 mm anterior, ±3.0 mm lateral, −8.4 mm ventral from the top of the skull).

### Drugs

Rats received saline or JHU37160 dihydrochloride (J60, Hello Bio, Princeton, NJ, 0.1 mg/kg, i.p.) 30 min prior to fear extinction, voltammetry, and/or locomotion procedures. J60 was dissolved immediately before use in sterile saline. 17β-estradiol benzoate (E2, Sigma Aldrich, St. Louis MO, 10 μg/250 g) was dissolved immediately before use in sesame oil (Sigma Aldrich, St. Louis MO) over low heat. Vehicle rats received 0.1 ml/250 g sesame oil. Cyclic injections commenced 4 to 5 days following OVX, whereby all rats received 1 subcutaneous injection of E2 every 4 days for 2 w before behavioral testing, as well as 30 min prior to extinction and FSCV testing.

### Fear Behavior

Females have been reported to have greater contextual fear conditioning relative to males [12, 33], so auditory fear conditioning was used in these experiments. Auditory fear conditioning (4 CS-US presentations; 10 sec, 80 dB, 2 KHz auditory CS terminating with a 1 sec, 0.8 mA foot shock US, 1 min ITI) took place in context A and auditory fear extinction (20 CS; 1 min ITI) occurred 24 h later in context B. The day after fear extinction, rats were again exposed to repeated presentations of the CS (between 5-20 CS, depending on experiment; 1 min ITI) in either the same context as where extinction took place (context B), or a different context than where extinction took place (context C). Rats were assigned to context B or context C based on freezing levels during extinction to avoid arbitrary differences in extinction levels between the contexts. Spontaneous recovery testing took place in context B 1 w later. Contexts A, B, and C differed in context shape, odor, lighting, and textures as previously described (Bouchet et al. 2018, and Supplementary Materials). All behavioral tests were recorded with overhead cameras and videos were later scored by both EthoVision XT (Leesburg, VA) and a human experimenter blind to treatment conditions of the animals.

### Locomotor Activity

Since conditioned freezing behavior can be influenced by locomotor activity as well as fear, locomotor activity was assessed during the 3 min period prior to the first CS presentation during each behavioral test using Noldus. Additionally, because females have been reported to express fear with high-velocity, darting behavior [34], high velocity movement throughout all behavioral tests was measured using Noldus. The effects of chemogenetic manipulations on locomotor activity was again assessed 24 h following the end of behavioral testing by placing rats into locomotor activity chambers (17″ × 17″ × 12″ L x W x H; Med Associates, Fairfax, VT) which use beam breaks to calculate the total distance traveled [32, 35].

### Fast scan cyclic voltammetry (FSCV)

SN neurons projecting to the DS contain DA [36], so FSCV was used to verify the anatomical specificity of the intersectional viral approach on DA release within DS subregions. Under urethane (1.5 g/kg, i.p.) anesthesia, a subset of rats used in behavioral experiments were placed in stereotaxic frames and 3-4 holes were drilled into the skull. A glass capillary-encased carbon fiber microelectrode (carbon fiber extending 120-150 μm beyond glass capillary) was lowered approximately 4.5-5.2 mm (from dura) into either the DMS (+0.5 mm anterior, ±1.8 mm lateral) or DLS (+0.5 mm anterior, ± 3.9 mm lateral from the top of the skull). A stimulating electrode (Plastics One, Roanoke, VA) was lowered 7.2-7.4 mm (from dura) into the SN (−5.4 mm anterior, ±3.0 mm lateral, from the top of the skull) and an Ag/AgCl reference electrode was implanted into the contralateral hemisphere just below the surface of the skull.

Voltammetric recordings were conducted by applying a triangular waveform (−0.4 V to 1.3 V; 400 V/s) which allowed for the detection of DA from cyclic voltammograms taken every 100 ms. To increase electrode sensitivity, the waveform was first applied at 60 Hz for ∼30 min, but was reduced to 10 Hz before experimentation. Data was collected in 15 s recordings taken 30 min after injections with 3 recordings taken every 5 min for each timepoint. Biphasic stimulation of the SN (24 rectangular pulses, 60 Hz, 300 μA) was delivered via the stimulating electrode 5 s into each recording. DA concentration was calculated from data collected during the recordings using linear regression and principle component regression as previously described [37].

### Tyrosine hydroxylase and cFos double immunohistochemistry

Male (n = 9) and cycling female rats were exposed to fear conditioning followed 24 h later by fear extinction. Female rats were grouped according to estrous phase during fear extinction (Met/Di n = 7; Pro/Est n = 11). Rats were trans-cardinally perfused with ice-cold saline and 4% paraformaldehyde 90 min after fear extinction. Double-label tyrosine hydroxylase (TH) and cFos immunohistochemistry and quantification occurred according to our previously published protocol [38].

### Statistical Analysis

Percent time spent freezing was calculated by averaging freezing data from the human scorer and scores from Noldus. Group differences in freezing and movement prior to the first CS in all contexts were analyzed with ANOVA. Group differences in freezing during fear conditioning were analyzed with repeated measures ANOVA. Freezing during fear extinction was collapsed into 10 blocks consisting of 2 CS each, and group differences were analyzed with repeated measures ANOVA. Freezing across all trials during fear renewal and spontaneous recovery were averaged and group differences were compared with multifactorial ANOVA. Group differences in locomotor activity were analyzed with multifactorial ANOVA. Changes in DA concentration relative to saline were analyzed with repeated measures ANOVA. Changes in DA concentration relative to vehicle were assessed using multifactorial ANOVA. Group differences in the % of TH-positive cells containing cFos were compared with ANOVA. The Shapiro-Wilk and Brown-Forsythe tests verified normality and equal variance of the data, respectively, prior to running ANOVAs. Bonferroni post-hoc analyses were performed when appropriate. Group differences were considered significant when p < 0.05.

## Results

### Sex and estrous phase during fear extinction modulate fear relapse and nigrostriatal DA

Here, we assessed the effects of sex and estrous phase during fear extinction on subsequent renewal and spontaneous recovery of fear (see Figure 1A for experimental design). No differences in freezing during any behavioral test were observed between females exposed to fear extinction during Met or Di and Pro or Est; therefore, females were assigned to Met/Di or Pro/Est groups based on estrous phase during extinction. Nine rats were excluded due to irregular or ambiguous estrous phases prior to fear extinction, resulting in final group sizes of n = 11 (Male/Same), n = 10 (Male/Different), n = 14 (Met/Di/Same), n = 7 (Met/Di/Different), n = 9 (Pro/Est/Same), n = 12 (Pro/Est/Different).

**Figure 1.**
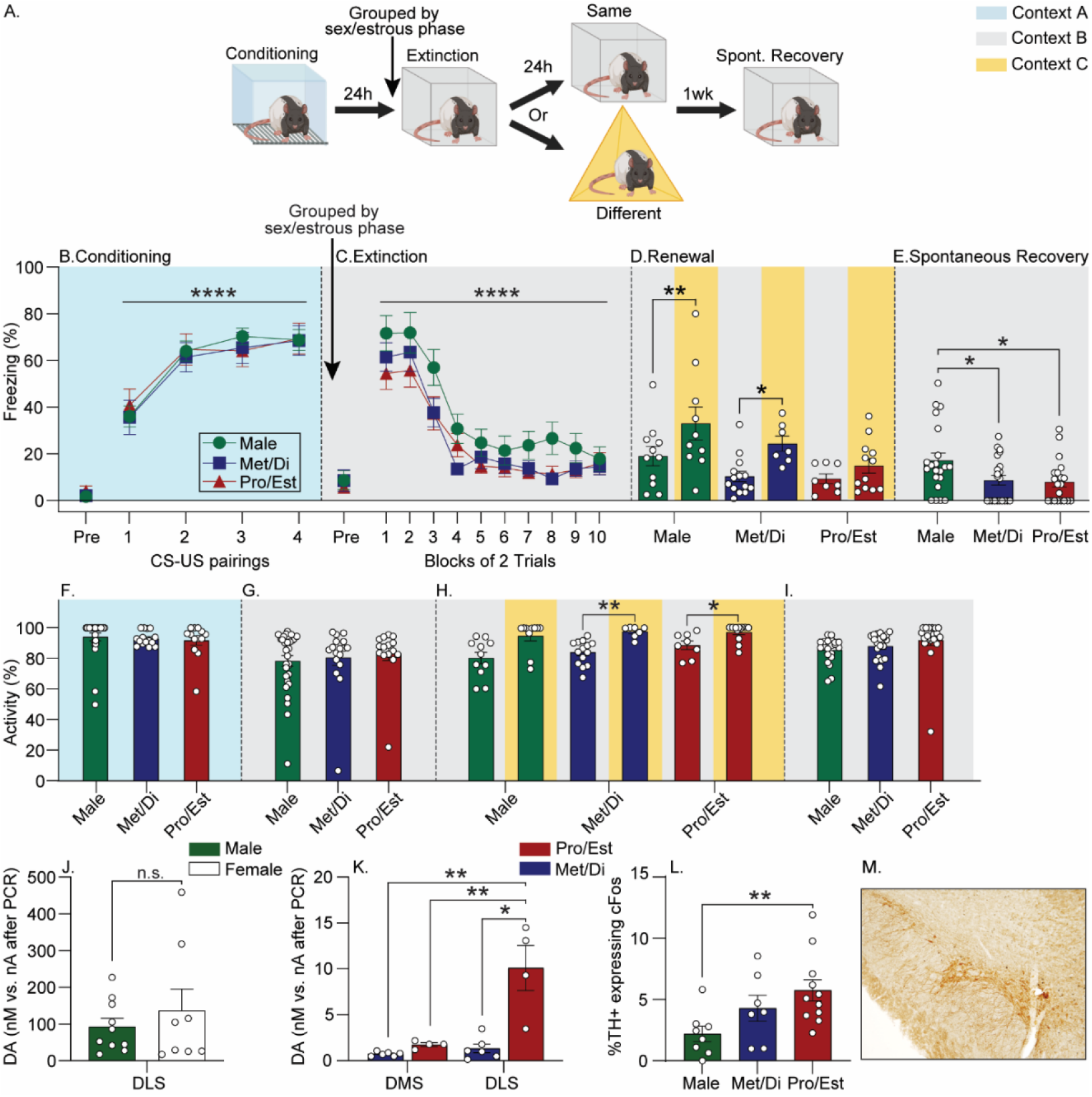
Effects of sex/estrous phase during auditory fear extinction training on fear relapse and nigrostriatal DA. A) Experimental design. Rats were exposed to auditory fear conditioning in context A (blue). The next day, rats were grouped by sex/estrous phase prior to fear extinction training in context B (grey) and 24 h later, re-exposed to the extinguished CS in either context B, or a novel context C to assess fear renewal. One w later, rats were placed back into context B and re-exposed to the CS to assess spontaneous recovery of fear. B-E) Levels of freezing during each behavioral session. F-I) Levels of locomotor activity prior to the first CS during each behavioral session. J) Effect of sex on electrically-evoked DA release in the DLS. K) Effect of female estrous phase on electrically-evoked DA release in the DMS and DLS. L) Percentage of tyrosine hydroxylase (TH)-positive neurons co-expressing cFos in the substantia nigra (SN) of rats exposed to fear extinction. M) Representative photomicrograph depicting double TH+ / cFos+ immunoreactivity in the SN. All freezing data represent group means ± SEM. *p<0.05; **p<0.01; ***p<0.001; ****p<0.0001, Bonferroni’s.

All rats acquired fear conditioning (F_(3,180)_ = 39.19; p < 0.0001; Figure 1B) and there were no differences in freezing between males and females subsequently assigned to Pro/Est or Met/Di groups (F_(2,60)_ = 0.25; p = 0.8; Figure 1B). All rats displayed within-session fear extinction (F_(9,540)_ = 69.31; p < 0.0001; Figure 1C) that was similar between sex/ extinction estrous phase groups (F_(2,60)_ = 1.94; p = 0.1; Figure 1C). Males and females in Met/Di during fear extinction displayed fear renewal in context C, whereas females in Pro/Est during fear extinction did not (main effect of sex/extinction estrous phase: F_(2,57)_ = 3.22; p < 0.05; main effect of context: F_(1, 57)_ = 7.26; p < 0.001; Figure 1D). There was no effect of sex/ extinction estrous phase on average freezing during the entire spontaneous recovery session (F_(2,60)_ = 0.56; p = 0.6; data not shown). However, when freezing in response to the first CS was analyzed separately to serve as a better representation of spontaneous recovery prior to the start of within-session extinction, we found that females froze less than males (F_(2,60)_ = 2.97, p < 0.05; Figure 1E). By this point, all females had been exposed to fear extinction or renewal sessions during Pro or Est, so it is likely that this sex difference was driven by high levels of ovarian hormones during either of these tests. Negligible high-velocity movement was detected during behavioral tests (range: 0.01 ± 0% - 0.54 ± 0.05%), so these data were not analyzed.

There were no sex or estrous phase differences in locomotor activity prior to the first CS during conditioning (F_(2,60)_ = 1.28; p = 0.3; Figure 1F), extinction (F_(2,60)_ = 1.19, p = 0.3; Figure 1G), or spontaneous recovery (F_(2,60)_ = 0.71, p= 0.5; Figure 1I). Rats placed into context C during renewal displayed increased locomotor activity compared to those placed into context B (F_(1,57)_ = 31.10; p < 0.0001; Figure 1H), but there was no effect of sex/estrous phase (F_(2,57)_ = 1.45; p = 0.2; Figure 1H).

We next evaluated how sex and estrous cycle impact the nigrostriatal DA pathway. We found no differences between females (n=8) and males (n = 10) in stimulus-evoked DA release in the DLS (F_(1,16)_ = 0.58; p < 0.45; Figure 1J). However, when estrous phases were analyzed separately, we found that females in Pro/Est (n=4) had greater stimulated DA release in the DLS compared to females in Met/Di (n=4; F_(1,6)_ = 5.884; p < 0.05; Figure 1K; Supplementary Figure 1) and this effect was observed to be greater in the DLS than the DMS (main effect of region: F_(1,6)_ = 6.577; p < 0.05; interaction between estrous phase and region: F_(1,6)_ = 5.884; p < 0.05; Figure 1G; Supplemental Figure 1). Female rats exposed to fear extinction during Pro/Est had a higher percentage of SN TH+ neurons containing cFos compared to males (F_(2, 23)_ = 4.37; p = 0.02; Figure 1L and 1M), indicating that estrous phase influences activity of SN DA neurons during fear extinction.

### Estradiol prior to fear extinction reduces fear relapse and increases stimulated DA release in the DLS

Here, we tested the hypothesis that E2 administration prior to fear extinction reduces fear relapse in OVX females (See Figure 2A for experimental design). One rat was excluded due to fatality from surgery. Group sizes were as follows: Vehicle/Same, n = 6, Vehicle/Different, n = 5; E2/Same, n = 5; E2/Different, n = 7. All rats acquired auditory fear conditioning (main effect of time: F_(3,63)_ = 14.27; p < 0.0001; Figure 2B), and freezing did not differ between drug groups (F_(1,21)_ = 2.55; p = 0.1; Figure 2B). All rats displayed within-session fear extinction (F_(9,189)_= 10.85; p < 0.0001; Figure 2C) that did not differ between drug groups (F_(1,21)_ = 2.47; p = 0.1; Figure 2C). Relative to vehicle, E2 prior to fear extinction reduced freezing during renewal (main effect of context: F_(1,19)_ = 4.03; p < 0.05; drug x context interaction: F_(1,19)_ = 4.80; p < 0.05; Figure 2D) and spontaneous recovery (F_(1,21)_ = 4.07; p < 0.05; Figure 2E). Again, very little high velocity movement was noted during all tests (range: 0.08 ± 0.16% - 0.84 ± 0.6%).

**Figure 2.**
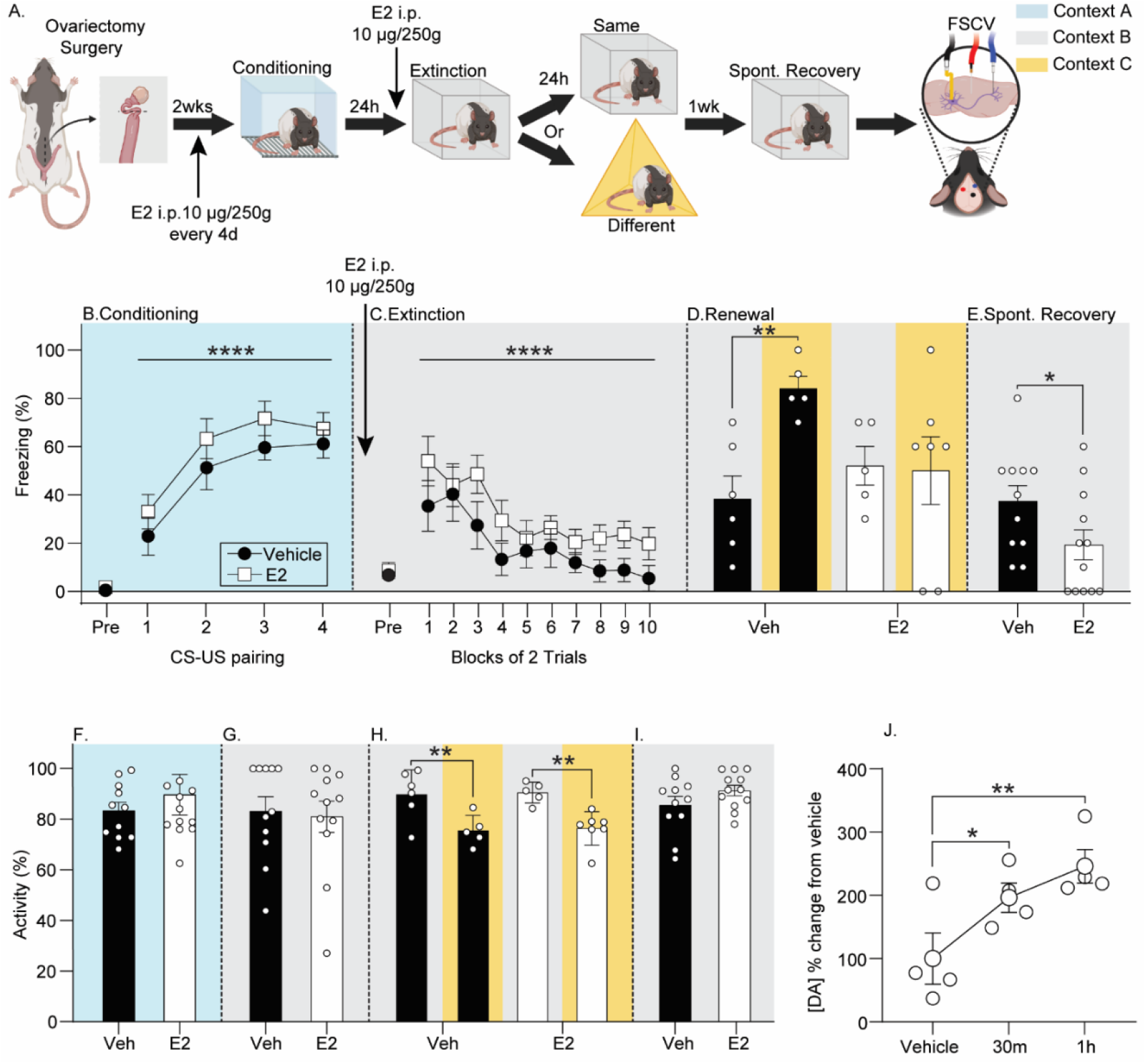
The effects of E2 administration during fear extinction training on fear relapse and dorsolateral striatum DA. A) Experimental design as described in Figure 1. Behavioral testing began 2 w following ovariectomy, during which time all rats received 10 μg/250g 17β-estradiol benzoate (E2) every 4 d. B-E) Levels of freezing during behavioral tests. Rats received either vehicle (Veh; 0.1 ml/250g) or E2 (10 μg/250g) administration 30 m prior to fear extinction training. F-I) Locomotor activity prior to the first CS during each behavioral test. J) Electrically-evoked DA release in the DLS after Vehicle administration or 30 m or 1 h after E2. All freezing data represent group means ± SEM. *p<0.05; **p<0.01; ***p<0.001; ****p<0.0001, Bonferroni’s.

Rats placed in context C during renewal testing displayed reduced locomotor activity prior to the first CS compared to rats placed into context B (F_(1,19)_ = 22.65; p < 0.0001; Figure 2H), but there was no effect of E2 (F_(1,19)_ = 0.07; p = 0.78; Figure 2H). No other group differences in locomotor activity prior to the first CS were noted (conditioning: F_(1,21)_ = 0.50; p = 0.5, Figure 2F; extinction: F_(1,21)_ = 0.06; p = 0.8, Figure 2G; spontaneous recovery: F_(1,21)_ = 2.23; p = 0.1, Figure 2I). Compared to vehicle, E2 increased stimulus-evoked DA release in the DLS of OVX rats (n = 4; F_(2,6)_ = 10.23; p < 0.01; Figure 2J; Supplementary Figure 2).

### Inhibition of the SN-DLS pathway during fear extinction restores fear renewal in females exposed to fear extinction during Pro/Est

Both stimulation of SN DA neurons [30] and increasing D1R signaling in the DLS at the time of fear extinction [31] reduce renewal in males, raising the possibility that potentiated DA release in the DLS observed during Pro/Est contributes to the acquisition of relapse-resistant fear extinction. Here, intersectional chemogenetics was used to inhibit the SN-DLS pathway during fear extinction in Pro/Est and Met/Di females and subsequent fear renewal was assessed (See Figure 3A for experimental design). After exclusion of rats with irregular estrous cycles (n = 2), fatalities from surgery (n = 7), and missed viral injection (n = 5), final group sizes were as follows: Met/Di/mCherry/Same, n = 8; Met/Di/mCherry/Different, n = 7; Met/Di/Gi-DREADD/Same, n = 9; Met/Di/Gi-DREADD/Different, n = 5; Pro/Est/mCherry/Same, n = 8; Pro/Est/mCherry/Different, n = 11; Pro/Est/Gi-DREADD/Same, n = 6; Pro/Est/Gi-DREADD/Different, n = 6.

**Figure 3.**
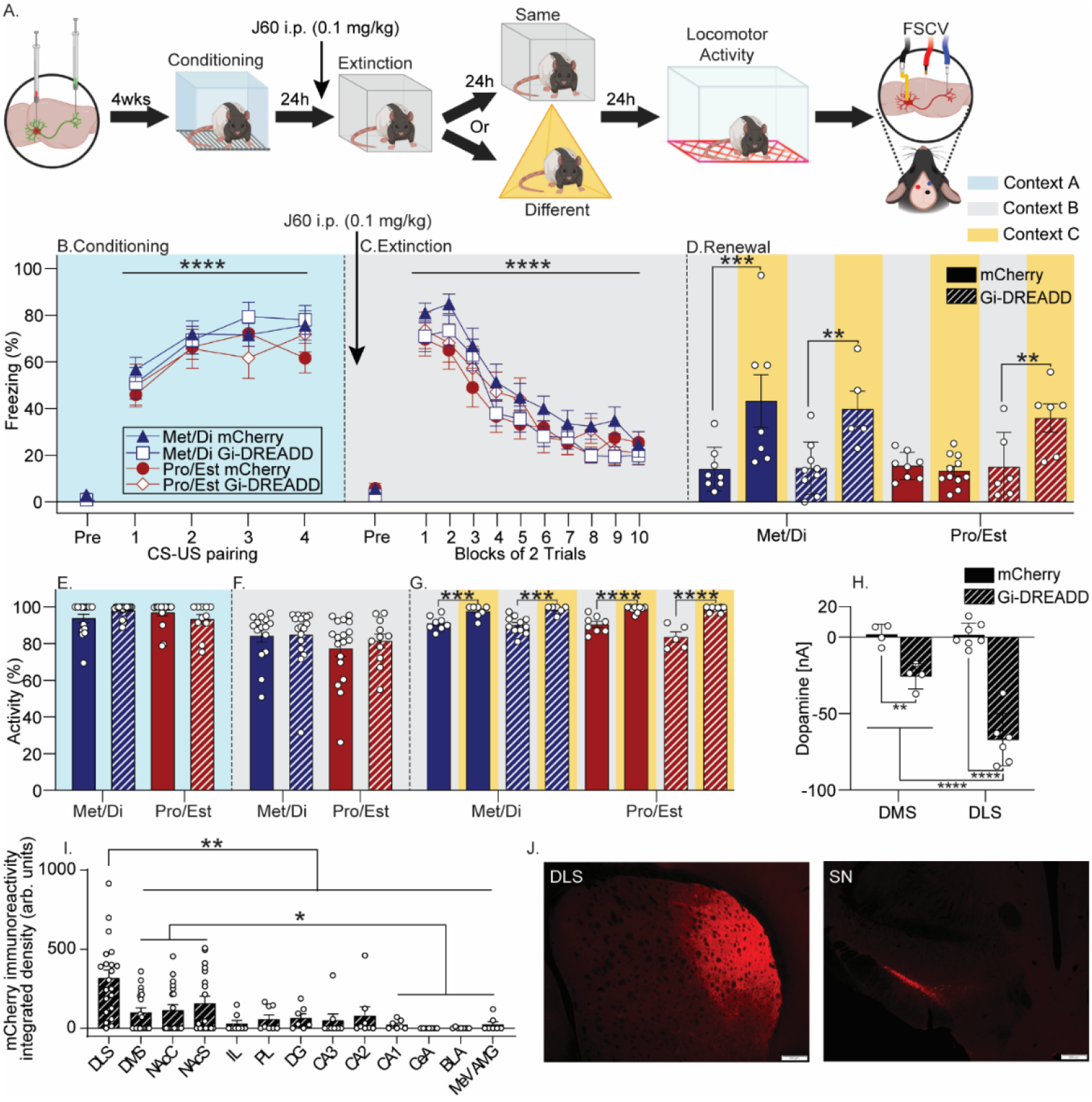
The effects of chemogenetic inhibition of substantia nigra (SN) neurons projecting to the dorsolateral striatum (DLS) during fear extinction on fear relapse in females. A) Experimental design as described in Figure 1. Females received injections of a retrogradeAAV encoding cre-recombinase into the DLS and either AAV-DIO-mCherry (mCherry) or an AAV containing a construct coding for a cre-recombinase-dependent Gi-coupled designer receptor exclusively activated by designer drug (Gi-DREADD) into the SN. Behavioral testing began 4 w after surgery. B-D) Levels of freezing during behavioral sessions. All rats received JHU37160 dihydrochloride (J60; 0.1 mg/kg, i.p.) 30 m prior to fear extinction training. E-G) Locomotor activity prior to the first CS during each behavioral test. H) Concentration of electrically-evoked DA release in the DLS following J60 administration. I) mCherry terminal immunoreactivity in striatal subregions and other brain regions implicated in fear extinction in rats expressing Gi-DREADD in the SN-DLS pathway analyzed with densitometry. K) Representative photomicrographs depicting mCherry in a rat injected with Gi-DREADD. Scale bar = 200μm. All freezing data represent group means ± SEM. *p<0.05; **p<0.01; ***p<0.001; ****p<0.0001, Bonferroni’s. Abbreviations: DLS, dorsolateral striatum; DMS, dorsomedial striatum; NAcC, nucleus accumbens core; NAcS, nucleus accumbens shell; IL, infralimbic cortex; PL, prelimbinc cortex; DG, dentate gyrus; CA3, cornu ammonis 3, CA2, cornu ammonis 2, CA1, cornu ammonis 1; CeA, central amygdala; BLA, basolateral amygdala; mAMG, medial amygdala.

All rats acquired auditory fear conditioning (F_(3,168)_ = 21.02; p < 0.0001; Figure 3B) that did not differ between virus (F_(1,56)_ = 0.01; p = 0.9; Figure 3B) or subsequently assigned estrous phase groups (F_(1,56)_ = 2.02; p = 0.2; Figure 3B). All rats displayed within-session fear extinction (F_(9,504)_ = 84; p < 0.0001; Figure 3C) that did not differ between estrous phase (F(1,56) = 0.84; p = 0.4; Figure 3C) or virus (F_(1,56)_ = 0.44; p = 0.5; Figure 3C). Inhibition of the SN-DLS pathway during fear extinction had no effect on fear renewal in Met/Di females, but restored fear renewal in Pro/Est females (main effect of estrous phase: F_(1,52)_ = 4.31; p < 0.05; main effect of context: F_(1,52)_ = 22.62; p < 0001; estrous phase x virus interaction: F_(1,52)_ = 5.44; p < 0.05; Figure 3D).

Negligible high velocity movement was again observed (range: 0.01 ± 0% - 1.50 ± 0.06%).

There were no group differences in locomotor activity prior to the first CS during conditioning (main effect of virus: F_(1,56)_ = 0.09; p = 0.8; Figure 3E; main effect of estrous phase: F_(1,56)_ = 0.04; p = 0.8; Figure 3E) or extinction (main effect of virus: F_(1,56)_ = 0.94; p = 0.3; Figure 3F; main effect of estrous phase: F_(1,56)_ = 0.46; p = 0.5; Figure 3F). Rats placed into context C during renewal displayed increased locomotor activity prior to the first CS compared to those placed into context B (F_(1,52)_ = 91.77; p < 0.0001; Figure 3G), but there was no effect of viral treatment (F_(1,52)_ = 1.29; p = 0.3; Figure 3G) or extinction estrous phase (F_(1,52)_ = 2.28; p = 0.1; Figure 3G). No effect of SN-DLS pathway inhibition on locomotor activity measured in locomotor activity chambers was found (Supplementary Figure 3).

FSCV was used to assess the ability of Gi-DREADD to suppress DA release in the DLS and DMS in a subset of rats expressing mCherry (n = 7) or Gi-DREADD (n = 6). J60 suppressed electrically-evoked DA release in the DLS and DMS in rats that received Gi-DREADD, relative to rats that received mCherry (F_(1,8)_ = 106.07; p < 0.0001; Figure 3H; Supplementary Figure 4). J60 elicited a larger suppression of DA release in the DLS than the DMS (virus x brain region interaction: F_(1,17)_ = 16.32; p < 0.001; Figure 3E; Supplementary Figure 4). DLS-projecting SN neurons branch and project to other regions [39]. Terminal mCherry immunoreactivity was highest in the DLS, but levels in other striatal regions were higher than in other brain areas (F_(17, 162)_ = 4.48; p < 0.0001; Figure 3I and 3J).

### Stimulation of the SN-DLS pathway during fear extinction reduces fear renewal in males

Here we used excitatory intersectional chemogenetics to test the hypothesis that SN-DLS pathway stimulation during fear extinction reduces relapse in males (experimental design shown in Figure 4A). After exclusion of fatalities from surgery (n = 5), statistical outliers (n = 3), and missed viral injections (n = 6), final group sizes were n = 8 (Male/mCherry/Same), n = 14 (Male/mCherry/Different), n = 14 (Male/Gq-DREADD/Same), n = 8 (Male/Gq-DREADD/Different).

**Figure 4.**
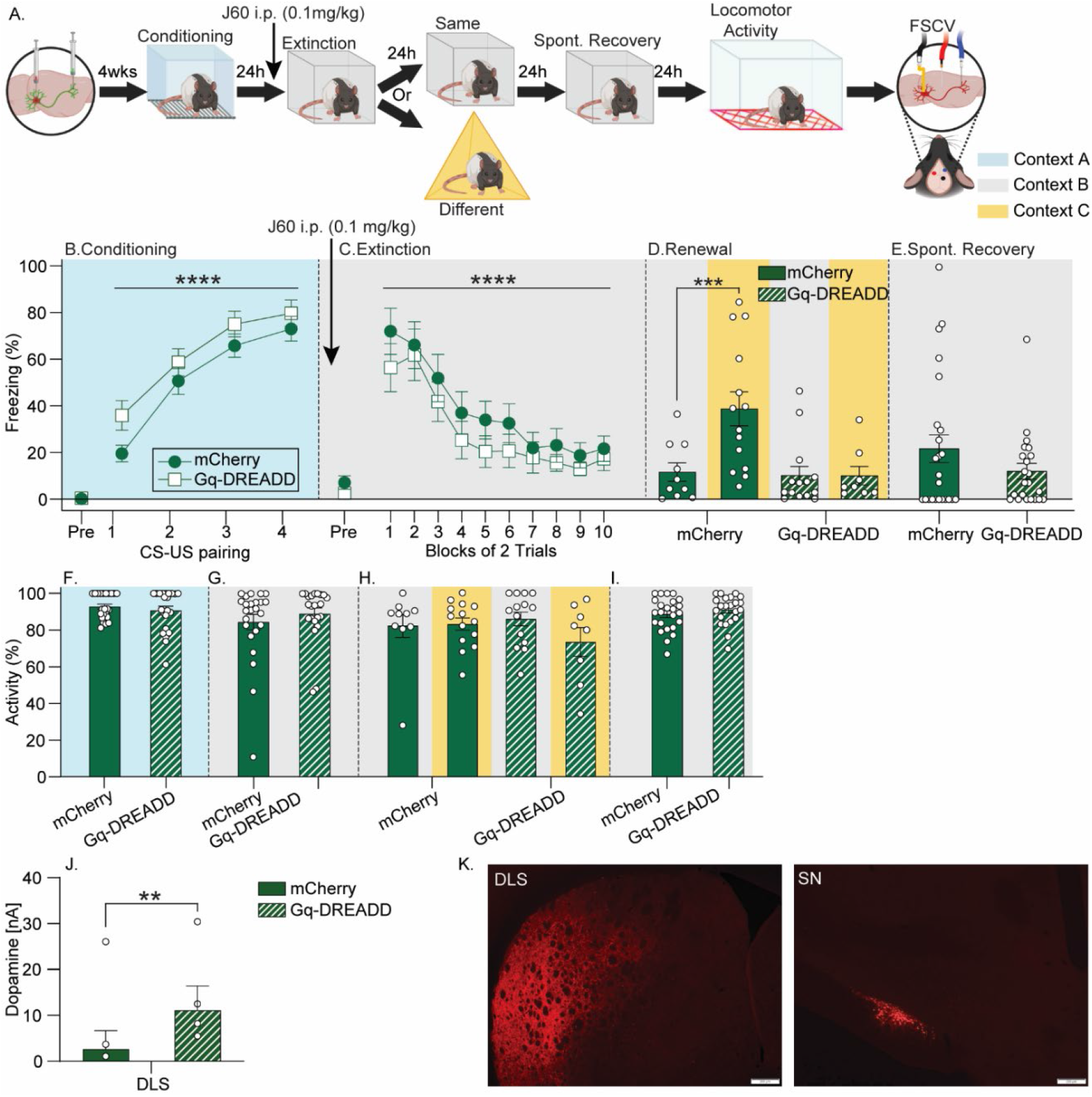
The effects of chemogenetic activation of substantia nigra (SN) neurons projecting to the dorsolateral striatum (DLS) during fear extinction on fear relapse in males. A) Experimental design as described in Figure 1. Male rats received injections of a retrograde AAV encoding cre-recombinase into the DLS and either AAV-DIO-mCherry (mCherry) or an AAV containing a construct coding for a cre-recombinase-dependent Gi-coupled designer receptor exclusively activated by designer drug (Gq-DREADD) into the SN. Behavioral testing began 4 w after surgery. B-E) Levels of freezing during behavioral sessions. All rats received JHU37160 dihydrochloride (J60; 0.1 mg/kg, i.p.) 30 m prior to fear extinction training. F-I) Locomotor activity prior to the first CS during each behavioral test. J) Concentration of electrically-evoked DA release in the DLS following J60 administration. K) Representative photomicrographs depicting mCherry in rats injected with Gq-DREADD. Scale bar = 200μm. All freezing data represent group means ± SEM. *p<0.05; **p<0.01; ***p<0.001; ****p<0.0001, Bonferroni’s.

All rats acquired auditory fear conditioning (F_(1,44)_ = 55.77; p < 0.0001; Figure 4B) that did not differ between virus (F_(1,44)_ = 2.63; p = 0.1; Figure 4B). All rats displayed within-session fear extinction (F_(1,44)_ = 20.43; p < 0.0001; Figure 4C) that did not differ between viral groups (F_(1,44)_ = 1.05; p = 0.3; Figure 4C). Rats that received mCherry displayed typical fear renewal, but stimulation of the SN-DLS pathway during fear extinction reduced renewal (main effect of virus: F_(1,40)_ = 6.69; p < 0.01; main effect of context: F_(1,40)_ = 4.85; p < 0.05; interaction between virus and context: F_(1,40)_ = 4.97; p < 0.05; Figure 4D). Levels of freezing during spontaneous recovery were not significantly different between groups (F_(1,44)_ = 1.85; p = 0.2; Figure 4E). Negligible high velocity movement was observed during all tests (range: 0.05 ± 0.01% - 0.93 ± 0.05%).

There were no group differences in locomotor activity prior to the first CS during conditioning (F_(1,44)_ = 0.67; p = 0.4; Figure 4F), extinction (F_(1,44)_ = 0.79; p = 0.4; Figure 4G), renewal (main effect of virus: F_(1,42)_ = 0.02; p = 0.9; main effect of context: F_(1,42)_ = 2.31; p = 0.1; Figure 4H), or spontaneous recovery (F_(1,44)_ = 0.90, p = 0.3; Figure 4I). No effect of SN-DLS pathway stimulation on locomotor activity measured in locomotor activity chambers was noted (Supplementary Figure 1B). J60 potentiated electrically-evoked DA release in the DLS of Gq-DREADD rats (n = 6) compared to mCherry (n = 6; F_(1,10)_ = 8.21; p < 0.01; Figure 4J; Supplementary Figure 5). Virus expressed robustly in the SN-DLS pathway (Figure 4K).

## Discussion

Here we report the novel finding that estrous phase during fear extinction modulates later fear relapse through a mechanism involving nigrostriatal DA. Females in Pro/Est during fear extinction had less fear renewal and spontaneous recovery than males and females in Met/Di during extinction. E2 seems to be the ovarian hormone important for mediating the observed effects of estrous cycle, as E2 administration in OVX females mimicked the effects of Pro/Est.

Relative to males, females in Pro/Est had greater activity of SN DA neurons during extinction, and both Pro/Est and E2 administration potentiated electrically-evoked DA release in the DLS. DLS DA is critical for ovarian hormone modulation of fear extinction and relapse, since SN-DLS pathway inhibition during fear extinction restored fear renewal in Pro/Est females and stimulating the SN-DLS pathway during extinction reduced fear renewal in males. These results provide new insight into how ovarian hormones modulate fear extinction in a relapse-resistant manner.

Results confirm our prior observation that estrous cycle during fear extinction training impacts later fear renewal [13]. This could potentially explain inconsistencies in the literature regarding sex differences in fear renewal. Binette et al. (2022) found that male rats have higher fear renewal after extinction than do females, whereas Schoenberg et al. (2024) reported no sex difference. However, whereas females in the Schoenberg et al. (2024) study were exposed to just one fear extinction training session, Binette et al. (2022) exposed cycling female rats to multiple fear extinction sessions, making it likely that many rats in the Benette et al. (2022) study were extinguished at least once while in Pro/Est. Future studies investigating sex differences in fear relapse should pay particular attention to the estrous phase or hormonal status of the females during the day of extinction.

In addition to impacting later fear renewal, we found that females with high levels of ovarian hormones during fear extinction (either through natural fluctuations or E2 administration) also have less spontaneous recovery compared to males and low-E2 females. This observation suggests that high levels of ovarian hormones during extinction could broadly reduce multiple forms of relapse in females, an observation with potential clinical implications.

The consensus of many reports suggests that ovarian hormones, particularly E2, promote the retention of fear extinction memories [5, 6, 8-11, 40, 41]. Notably, this effect was not observed here or in our prior work [13]. One explanation for this discrepancy could be differences in the strength of the fear conditioning memory between studies. We specifically use conditioning and extinction conditions that elicit fear renewal. This necessitates low levels of freezing in the “same” context, which could obscure potential group differences in fear extinction retention in this context. Indeed, most of the prior work reporting that estrous cycle at the time of fear extinction impacts extinction retention used a greater number of CS-US pairings during conditioning than we use in the current studies.

Understanding the effects of estrous cycle during fear extinction on fear extinction retention is important because it can provide insight into how ovarian hormones during extinction impact later relapse. If improved extinction retention in Pro/Est groups was obscured by a floor effect in the current studies, the reduction in relapse observed in Pro/Est females could nonetheless be a product of improved fear extinction retention. Alternatively, ovarian hormones during fear extinction could impact fear relapse through a mechanism independent of enhanced fear extinction memory, *per se*. Supporting this possibility, the reduced fear renewal in female, compared to male, rats reported by Binette et al. (2022) similarly occurred in the absence of improved fear extinction retention in females. Moreover, both stimulation of the SN-DLS pathway (Figure 4D) and increasing D1R signaling in the DLS during fear extinction [31] reduce fear renewal without enhancing extinction memory. Since high levels of ovarian hormones enhance SN DA neural activity during fear extinction, potentiate electrically-evoked DA release in the DLS, and depend on SN-DLS pathway activity during fear extinction, it seems possible that ovarian hormones could recruit a mechanism involving DA in the DLS that renders fear extinction memory resistant to relapse independently from the strength of the fear extinction memory.

We observed that estrous cycle impacts the nigrostriatal DA pathway at both the cell body and terminal regions. Although the percentage of SN DA neurons expressing cFos after fear extinction was very low, it was nonetheless greater in females exposed to fear extinction during Pro/Est compared to Met/Di and males. Since ovarian hormones are thought to modulate DA transmission through DA terminal disinhibition rather than acting directly at the cell body [42, 43], greater extinction-induced SN DA activity observed during Pro/Est could be mediated by feedback loops through the SN reticularis [39]. Within the DS, we found that estrous phase had a

more pronounced effect on DA release in the DLS than the DMS, suggesting that ovarian hormones can influence DA transmission differently in different DA terminal regions. Together, the data support the possibility that high levels of ovarian hormones during fear extinction could reduce relapse by potentiating DA release in the DLS, but they do not rule out a potential contribution from DA in other striatal regions, which is also known to be potentiated by high levels of ovarian hormones.

The observation that inhibition of the SN-DLS pathway during fear extinction did not impact fear extinction acquisition, retention, or renewal in Met/Di females is consistent with our recent observation that temporary inhibition of the DLS during fear extinction has no impact on fear extinction or renewal in male rats [31]. These data suggest that the SN-DLS pathway may not contribute to “normal” fear extinction or relapse. However, SN-DLS pathway inhibition during fear extinction restored renewal in females exposed to fear extinction during Pro/Est.

Thus, when it is recruited during fear extinction by the presence of ovarian hormones, heightened activity of the SN-DLS pathway renders fear extinction memory resistant to relapse. In fact, stimulation of the SN-DLS pathway during fear extinction seems to be sufficient to reduce renewal in the absence of high levels of ovarian hormones, as stimulation of the SN-DLS pathway (Figure 4D) or a D1R agonist injected into the DLS [31] during fear extinction both reduce renewal in males. These data suggest that targeting SN-DLS pathway activity during fear extinction could be a means to reduce relapse in both sexes.

We assessed the anatomical specificity of the intersectional approach to target the SN-DLS pathway. The highest levels of terminal mCherry were observed in the DLS, consistent with the large suppression of electrically-evoked DA caused by J60 in rats expressing Gi-DREADDs. However, some mCherry was observed in other striatal regions, and J60 also suppressed DA release in the DMS, albeit to a lesser degree than the DLS. The potential role of alterations in DA transmission in other striatal regions contributing to the effects of estrous cycle or chemogenetic manipulations; therefore, cannot be entirely ruled out. These data also have important implications for other studies utilizing intersectional chemogenetics to target “projection-defined” DA pathways.

Interestingly, the current results add to growing evidence suggesting that females, compared to males, preferentially recruit DLS circuits to guide behavior. Formation of habitual responding during operant training involves a shift from DMS- to DLS-control of operant behavior [44, 45], and females form habits earlier during operant training than do males [46, 47]. Female rats given the opportunity to press a lever to escape from electric shock do so with the DLS, whereas males use the DMS to accomplish the identical instrumental task [48]. When provided with a running wheel in their cages, both sexes of rats will engage in robust wheel running behavior. However, females have a preference to use the DLS to govern the acquisition of wheel running, whereas males prefer the DMS [49]. Here, we find that females also recruit DLS circuits during fear extinction, a task not typically associated with the DS, and that this is influenced by ovarian hormones.

In summary, high levels of ovarian hormones present during fear extinction render fear extinction resistant to relapse through a nigrostriatal pathway. Future work is required to understand how SN-DLS pathway activity during extinction reduces later relapse. Regardless of the mechanism, the SN-DLS pathway should be considered as a potential target for the reduction of fear relapse after extinction in both sexes.

## Notes

### Competing Interest Statement

The authors have declared no competing interest.

