## Supplemental Materials and Methods for "Estrous phase during fear extinction modulates fear relapse through a nigrostriatal dopamine pathway"

**Supplementary Figures**

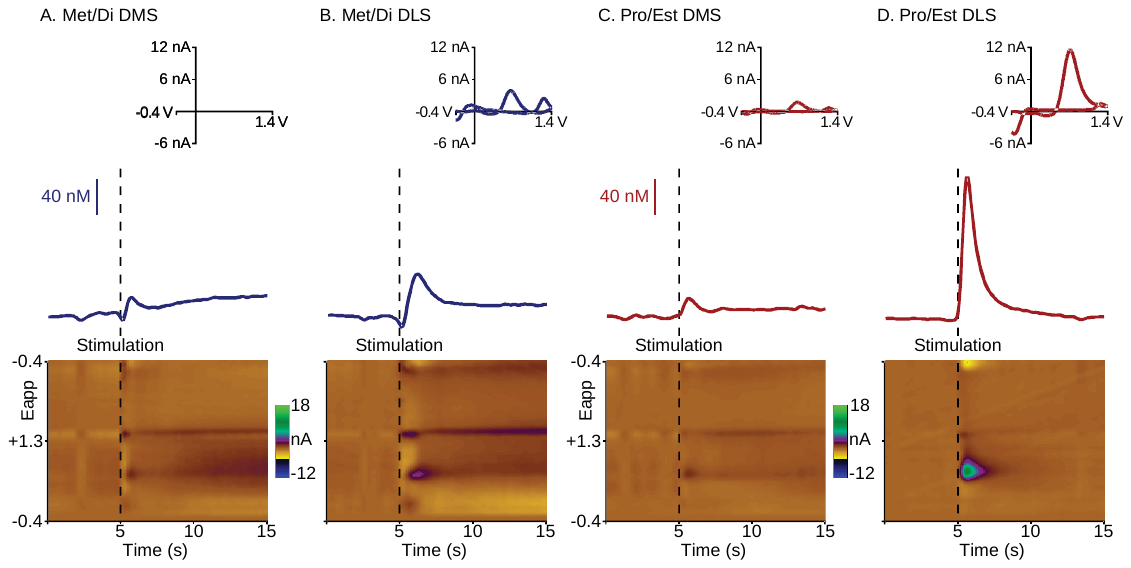

Supplementary Figure 1. Electrically-evoked dopamine release in dorsal striatal subregions during different phases of the estrous cycle (see Figure 1). Representative examples are shown of measurements taken in the DMS **(A)** and DLS **(B)** of a single rat during the Metestrus or Diestrus (Met/Di) phase of the estrous cycle, as well as the DMS **(C)** and DLS **(D)** of a single rat during Proestrus or Estrus (Pro/Est). **(A, B, C and D)** The effect of electrical stimulation on dopamine release is shown as color plots (Bottom) and corresponding concentration traces (Middle) with cyclic voltammograms (Top). Each set of plots represents data collected during a single recording. Color plots depict time (seconds; x-axis), scan potential applied to the electrode (Eapp [V]; y-axis), and voltammetric current (nA; z-axis). Electrical stimulation (24 pulses, 60 Hz, 300 μA, 2 ms/phase, biphasic) of the substantia nigra is indicated by the black dashed lines. Following stimulation, a dopamine release event is observed in the color plots at the oxidative potential for dopamine (+0.6 V). The corresponding concentration traces illustrate the concentration of dopamine (nM; y-axis) across time (seconds; x-axis). Concentration (nM) was converted from current (nA) using a calibration factor. Cyclic voltammograms represent the peak concentration seen in the concentration traces following electrical stimulation and verify the signal as dopamine. Cyclic voltammograms are plotted as current (nA; y-axis) as a function of applied potential (V; x-axis).

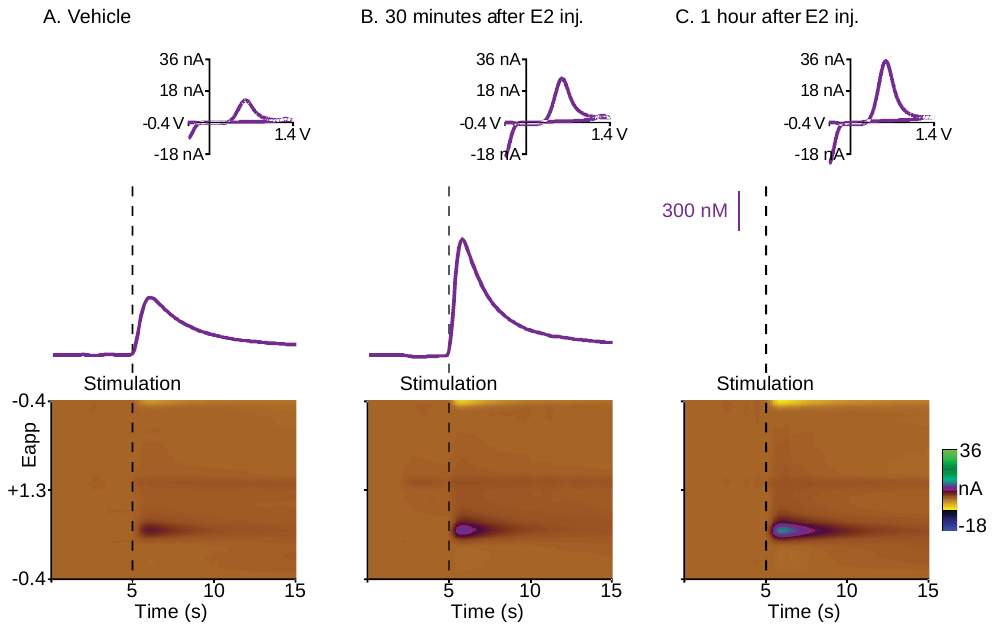

Supplementary Figure 2. Effect of estradiol (E2) administration on electrically-evoked dopamine release in the dorsolateral striatum (DLS) in ovariectomized (OVX) female rats (see Figure 2). Recordings were taken 30 min after vehicle administration to establish a baseline response to electrical stimulation. E2 was then administered and recordings were taken 30 min and 1 h after injection. Representative examples are shown of dopamine measured in the DLS of a single rat 30 min after vehicle injection **(A)**, 30 min after E2 **(B)**, and 1 h after E2 **(C)**. During each recording, the SN was electrically stimulated (24 pulses, 60 Hz, 300 μA, 2 ms/phase, biphasic) after 5 s and subsequent dopamine release events were recorded. **(A-C)** The effect of electrical stimulation on dopamine release is shown as color plots (Bottom) and corresponding concentration traces (Middle) with cyclic voltammograms (Top). Each set of plots represents data collected during a single recording. Stimulation is indicated in the color plots and concentration traces by the black dashed lines. See Figure S1 for a description of the representative voltammetric plots.

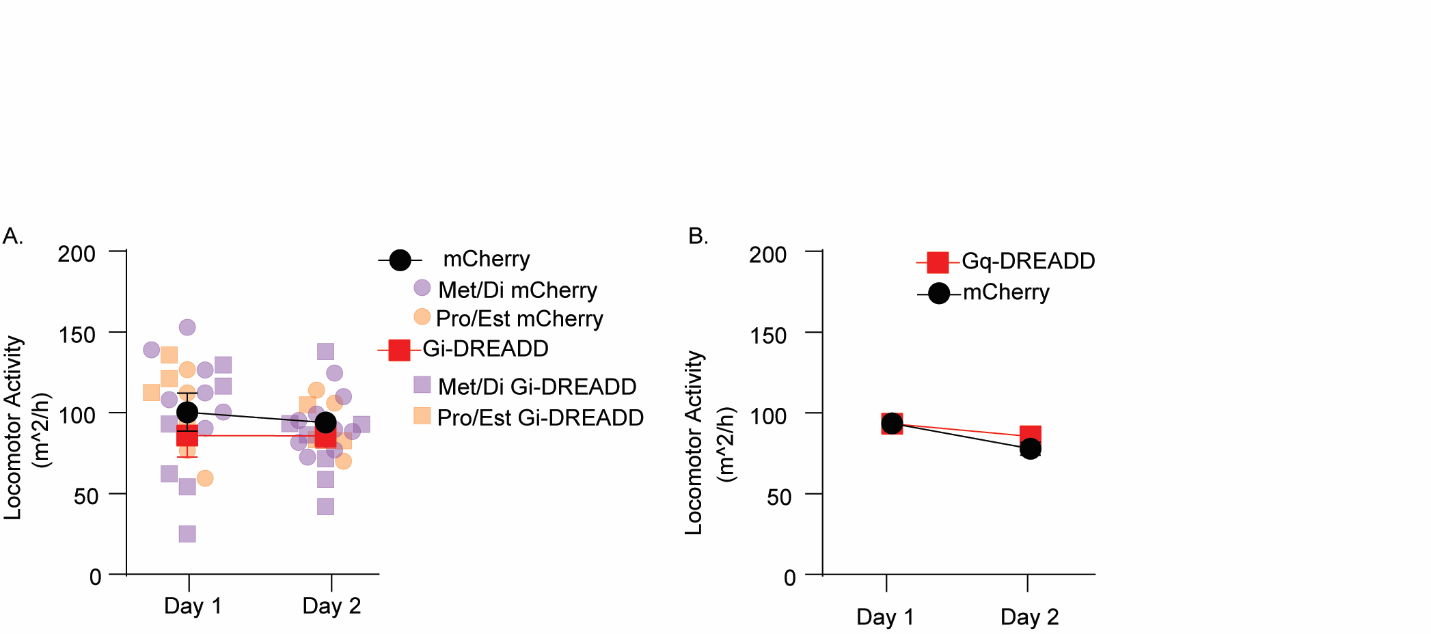

Supplementary Figure 3. Effects of chemogenetic manipulations on locomotor activity. **(A)** Female rats expressing cre-recombinase-dependent mCherry or Gi-DREADD in the substantia nigra – to – dorsolateral striatum (SN-DLS) pathway were placed into Med Associates locomotor activity chambers for 1 h per day, for 2 consecutive days. DREADD ligand JHU37160 dihydrochloride (J60; 0.1 mg/kg, i.p.) was administered 30 min prior to placement into locomotor activity chambers on day 1 only. Estrous phases were identified with vaginal lavage prior to behavioral testing on both days. Locomotor activity decreased over time (F_(1,57)_ = 15.51; p < 0.02), but no difference between mCherry or Gi-DREADD was observed (F_(1,57)_ = 0.51; p = 0.47). Data for individual rats in metestrus or diestrus (Met/Di) or proestrus or estrus (Pro/Est) are depicted on days 1 and 2. **(B)** Male rats expressing mCherry or Gq-DREADD in the SN-DLS pathway were placed into Med Associates locomotor activity chambers for 1 h per day, for 2 consecutive days. DREADD ligand J60 (0.01 mg/kg i.p.) was administered 30 min prior to placement into locomotor activity chambers on day 1 only. Locomotor activity decreased over time (F_(1,44)_ = 15.4; p < 0.03), but no difference between mCherry or Gi-DREADD was observed (F_(1,44)_ = 0.33; p = 0.56). All data represent groups means ±SEM.

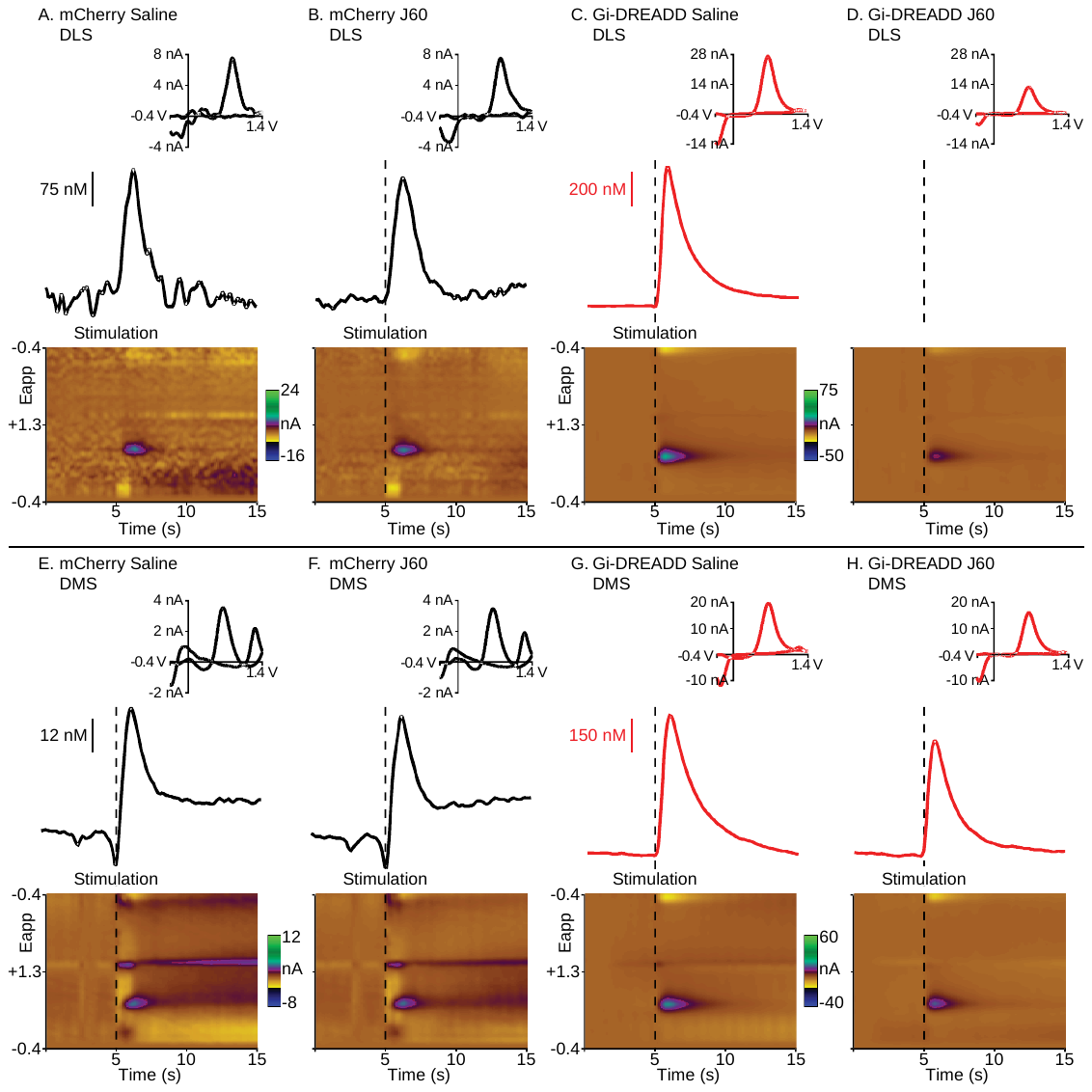

Supplementary Figure 4. Effect of JHU37160 dihydrochloride (J60; 0.1 mg/kg, i.p.) administration on electrically-evoked dopamine release in dorsal striatal subregions of rats expressing either mCherry control virus or Gi-DREADD virus in the substantia nigra (SN) – to – dorsolateral striatum (DLS) pathway (see Figure 3). Recordings were taken 30 min after vehicle injection to establish a baseline dopamine response to electrical stimulation. J60 was then given, and recordings were taken 30 min after injection. Representative fast scan cyclic voltammetry (FSCV) data are shown of recordings taken in the DLS of a single mCherry virus control rat 30 min after vehicle injection **(A)** and 30 min after J60 injection **(B)**, as well as recordings taken in the DLS of a single Gi-DREADD virus rat 30 min after vehicle injection **(C)** and 30 min after J60 injection **(C)**. Recordings were taken in the DMS to determine the specificity of the intersectional dual virus approach used to target the SN-DLS pathway. Representative examples are shown of recordings taken in the DMS of a single mCherry control rat 30 min after vehicle **(E)** and 30 min after J60 **(F)**, as well as recordings taken in the DMS of a single Gi-DREADD virus rat 30 min after vehicle **(G)** and 30 min after J60 **(H)**. During each recording, the SN was electrically stimulated (24 pulses, 60 Hz, 300 μA, 2 ms/phase, biphasic) after 5 s and subsequent dopamine release events were recorded. **(A-H)** The effect of electrical stimulation on dopamine release is shown as color plots (Bottom) and corresponding concentration traces (Middle) with cyclic voltammograms (Top). Each set of plots represents data collected during a single recording. Stimulation is indicated in the color plots and concentration traces by the black dashed lines. See Figure S1 for a description of the representative voltammetric plots.

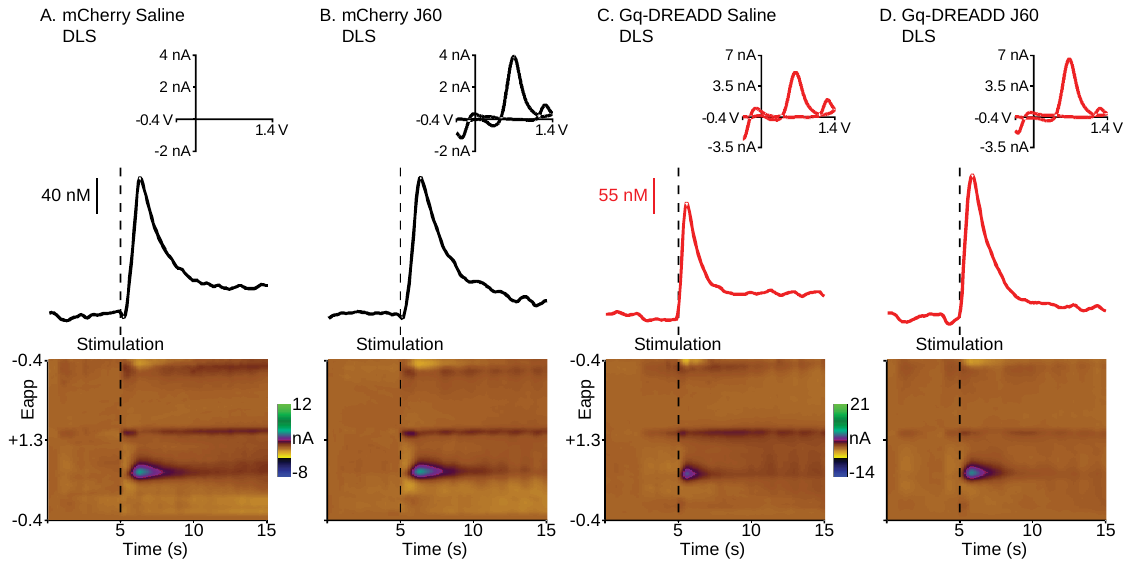

Supplementary Figure 5. Effect of JHU37160 dihydrochloride (J60; 0.1 mg/kg, i.p.) administration on electrically-evoked dopamine release in the dorsolateral striatum (DLS) of rats expressing either mCherry control virus or Gq-DREADD virus in the substantia nigra (SN) – to – dorsolateral striatum (DLS) pathway (see Figure 4). Recordings were taken 30 min after vehicle injection to establish a baseline dopamine response to electrical stimulation. J60 was then given, and recordings were taken 30 min after injection. Representative fast scan cyclic voltammetry (FSCV) data are shown of recordings taken in the DLS of a single mCherry control rat 30 min after vehicle injection **(A)** and 30 min after J60 injection **(B)**, as well as recordings taken in the DLS of a single Gq-DREADD virus rat 30 min after vehicle injection **(C)** and 30 min after J60 injection **(C)**. During each recording, the SN was electrically stimulated (24 pulses, 60 Hz, 300 μA, 2 ms/phase, biphasic) after 5 s and subsequent dopamine release events were recorded. (**A-D)** The effect of electrical stimulation on dopamine release is shown as color plots (Bottom) and corresponding concentration traces (Middle) with cyclic voltammograms (Top). Each set of plots represents data collected during a single recording. Stimulation is indicated in the color plots and concentration traces by the black dashed lines. See Figure S1 for a description of the representative voltammetric plots.

Supplementary Materials and Methods

Estrous phase identification

Vaginal epithelial cells were collected using a sterile, blunt-tipped eye dropper filled with ~0.5 mL sterile-filtered 0.2% PBS-Brij solution (Brij 35 Solution 30%; Sigma Aldrich, B4184). The vagina was gently flushed, and the collected fluid was transferred onto a microscope slide for visualization under a 20X objective lens (Olympus BX53).

Ovariectomy and Estrogen Administration

Female rats were bilaterally ovariectomized (n=­23) as previously described (Tanner et al., 2023). Briefly, under ketamine (75.0 mg/kg i.p.) and medetomidine (0.5 mg/kg i.p.) anesthesia, a dorsal incision was made in the skin followed by bilateral incisions through the muscle wall. Ovaries were located, the uterine horns were ligated with absorbable Vicryl sutures (4-0, FS-2), and the ovaries were removed. The muscle and skin incisions were closed with absorbable sutures. Bupivacaine (2mg/kg) was applied at the sight of muscle incision prior to skin closure. Injections of carprofen (5mg/kg s.c.) and penicillin G (22,000 IU/rat s.c.) were administered at induction and every 24 h for 72 h after surgery. Rats recovered for 2 weeks prior to experimentation. Vaginal lavage after OVX or Sham surgery verified lack of cycling in 100% of OVX and 0% of Sham rats.

mCherry Densitometry

Immunohistochemistry (IHC) was performed on brain sections containing (from rostral to caudal) prefrontal cortex (3.7mm to 1.70mm rostral from Bregma), striatum (1.6mm to 0.2mm rostral from Bregma), hippocampus/amygdala (−2.12mm to −4.52mm caudal from Bregma) and substantia nigra (-4.8mm to -6.04mm caudal from Bregma). Sections were rinsed 3 times for 10 minutes using 0.01M phosphate buffed saline (PBS), followed by an overnight incubation at room temperature in 5% blocking solution containing 0.3% Triton X, 0.01M PBS, and 5% normal goat serum (NGS). Sections were then rinsed 3 times for 10 minutes in 0.01M PBS, the sections were placed in 3% blocking solution of 0.01M PBS, 5% NGS, and rabbit anti-mCherry at 1:50000 for an overnight incubation at room temperature. Sections were then rinsed 3 times for 10 minutes in 0.01M PBS, then sections were incubated in 3% blocking solution containing 0.01M PBS, 5% NGS, and Alexa Fluor goat anti-rabbit 594 at 1:250 for 4 hours at room temperature. Sections were then rinsed 3 times for 10 minutes in 0.01M PBS, and tissue was mounted on slides using deionized water and left to airdry slightly before cover slipping with Prolong Gold with dapi. Images through SN projection sites implicated in fear extinction were captured from at least 8 hemispheres per brain region per animal, and intensity of mCherry signal above background (densitometry) was calculated as previously described [50].

Behavioral Procedures

Fear conditioning, extinction, and renewal tests were separated by 24 h, while spontaneous recovery took place 1 week following fear extinction, and occurred in distinct contexts (A, B, C design). Schematics depicting experimental designs are shown in Figures 1A, 2A, 3A and 4A. Freezing was defined as the absence of all movement except that required for respiration and was used as a measure of fear in all behavioral tests. Behavior was recorded with overhead cameras and videos were later scored both with EthoVision XT (Leesburg, VA) and by a human experimenter blind to treatment conditions of the animals. EthoVision was used to quantify locomotor activity prior to the first CS presentation during each behavioral test.

Rats were transported in their home cages to a behavioral testing room to undergo fear conditioning. Rats were placed into custom, rectangular conditioning chambers (context A; 20” W x 10” D x 12” H) with a shock grid floor (Coulbourn Instruments, Allentown, PA). Conditioning chambers were contained within individual sound-attenuating cabinets illuminated by red lights. Rats were allowed 3 min to explore the context, after which 4 auditory CS (10 sec, 80 dB, 2 KHz), each co-terminating with a 1 sec, 0.8-mA foot shock US were delivered on a 1 min inter-trial interval. Auditory stimuli and foot shocks were delivered through Coulbourn tone generators and shock scramblers controlled via Noldus EthoVision XT software through a custom interface. Rats remained in the conditioning chamber for 1 min after the last shock before being transported back to the housing room. Conditioning chambers were cleaned with water between rats.

During auditory fear extinction in context B, rats were placed into either a custom Plexiglas chamber that was either rectangular (15” W x 15” D x 20” H) with a textured floor or a triangular (15” sides x 20” H) with a smooth floor. Both chambers were counterbalanced so that half the rats were exposed to fear extinction in the rectangular chambers and the other half in the triangular chambers. Fear extinction took place in the same sound-attenuating cabinets as conditioning. Cabinets contained vanilla scent and were illuminated with bright white lights, while a fan inside the cabinet provided ventilation and background noise. Rats were transported to the sound-attenuating cabinets in their assigned chambers. After a 3 min exploratory period, the auditory CS was presented 20 times (1 min ITI) in the absence of the foot shock US. Rats were removed from context B 1 min after the last auditory CS presentation and returned to their homecages in the housing room. context B chambers were cleaned with 10% ethanol between tests.

During fear renewal testing, half of the rats were re-exposed to the auditory CS in the same context that fear extinction took place (context B; Same) while the other half were re-exposed to the auditory CS in a different context than where fear extinction took place (context C; Different), so that rats that underwent fear extinction in the rectangular Plexiglas chamber were now placed into the triangular chamber, and vice versa. Rats in context B chambers were transported to the sound attenuating cabinets with identical treatment conditions as context B during fear extinction. Rats in context C chambers were transported to the sound attenuating cabinets consisting of a raspberry scent and dimly lit by a lamp located outside of the cabinets. After a 3 min exploration period, the auditory CS was presented 20 times (1 min ITI) in the absence of the foot shock US. Rats were removed from context B or context C 1 min after the last auditory CS presentation and returned to their home cages in the housing room. Context C chambers were cleaned with 1% acetic acid between tests. Experimenters transporting and handling rats also differed between contexts, such that unique experimenters were used for each context. In the experiment in which E2 was administered to OVX rats, 5 CS presentations were used during fear renewal testing to avoid floor effects during spontaneous recovery.

During spontaneous recovery, rats were transported in context B chambers to the sound attenuating cabinets with identical treatment conditions as fear extinction. After a 3 min exploration period, the auditory CS was presented 20 times (1 min ITI) in the absence of the foot shock US. Rats were removed from context B 1 min after the last auditory CS presentation and returned to their home cages in the housing room. Context B chambers were cleaned with 10% ethanol between tests.
